# Structural insights into ligand efficacy and activation of the glucagon receptor

**DOI:** 10.1101/660837

**Authors:** Daniel Hilger, Kaavya Krishna Kumar, Hongli Hu, Mie Fabricius Pedersen, Lise Giehm, Jesper Mosolff Mathiesen, Georgios Skiniotis, Brian K. Kobilka

**Author notes:** contributed equally (Daniel Hilger, Kaavya Krishna Kumar, Hongli Hu).

## Abstract

The glucagon receptor family comprises Class B G protein-coupled receptors (GPCRs) that play a crucial role in regulating blood sugar levels. Receptors of this family represent important therapeutic targets for the treatment of diabetes and obesity. Despite intensive structural studies, we only have a poor understanding of the mechanism of peptide hormone-induced Class B receptor activation. This process involves the formation of a sharp kink in transmembrane helix 6 that moves out to allow formation of the nucleotide-free G protein complex. Here, we present the cryo-EM structure of the glucagon receptor (GCGR), a prototypical Class B GPCR, in complex with an engineered soluble glucagon derivative and the heterotrimeric G-protein, G_s_. Comparison with the previously determined crystal structures of GCGR bound to a partial agonist reveals a structural framework to explain the molecular basis of ligand efficacy that is further supported by mutagenesis data.

## Main Text

The peptide hormone glucagon is released from the α cells of the pancreas in response to low blood glucose level^1^. Glucagon binds and activates the glucagon receptor (GCGR), a member of the Class B family of GPCRs^2,3^. Upon activation, GCGR potentiates signaling predominantly by interacting with the adenylyl cyclase stimulatory G protein, G_s_, and upregulating cAMP production^4^. This ultimately leads to enhanced glycogenolysis and gluconeogenesis and an increased hepatic glucose output^5^. Due to its fundamental role in glucose homeostasis, GCGR represents an important therapeutic target for the treatment of severe hypoglycemia in diabetic patients^6^. Furthermore, modulation of GCGR signaling has been shown to have therapeutic potential for obesity and type 2 diabetes therapies^7^. Structural studies on GCGR have provided important insights into negative allosteric modulation of the receptor^8,9^. However, the molecular basis of receptor activation by the endogenous hormone glucagon is not well understood. Although, the recently determined partial agonist-bound GCGR structure offers an overview into peptide-binding, it does not reveal the allosteric link between peptide binding and activation^10^. Moreover, the partial-agonist peptide lacks residues that are present in glucagon and have been shown to be important determinants of efficacy^11^. Structures of other Class B receptor:G protein complexes have shown, unlike in Class A receptor complexes, the formation of a complete break in the TM6 helix that accommodates the α5 helix of Gα_s_^12-16^. Since the partial agonist-bound GCGR structure was solved in the absence of G protein, the fully active conformation of the receptor with a “kinked” TM6 was not captured.

In order to determine the structure of the GCGR signaling complex, we engineered a glucagon analogue (ZP3780), that shows significantly improved solubility and stability in aqueous solutions in comparison to the native glucagon peptide that is prone to rapidly form fibrillar aggregates at neutral pH (Suppl. Fig. 1, Suppl. Table 1, 2)^17^. Taking advantage of our previous efforts in designing glucagon analogues^18^, ZP3780 was engineered to act as a full agonist by retaining the native sequence in the N-terminus of the peptide, important for ligand efficiency^19^, while improving solubility and stability by iteratively substituting amino acids in the C-terminus to change the peptide isoelectric point (pI) (Suppl. Fig. 1). The resulting glucagon analogue ZP3780 includes four C-terminal mutations that only slightly reduce the affinity in comparison to the wild-type glucagon (Suppl. Fig. 1). Activation of the receptor with ZP3780 enabled the formation of a GCGR:G_s_ complex that was stable enough for cryo-electron microscopy (cryo-EM) imaging yielding a final density map at a nominal resolution of 3.1Å (Fig. 1a, Suppl. Fig. 2).

**Fig. 1:**
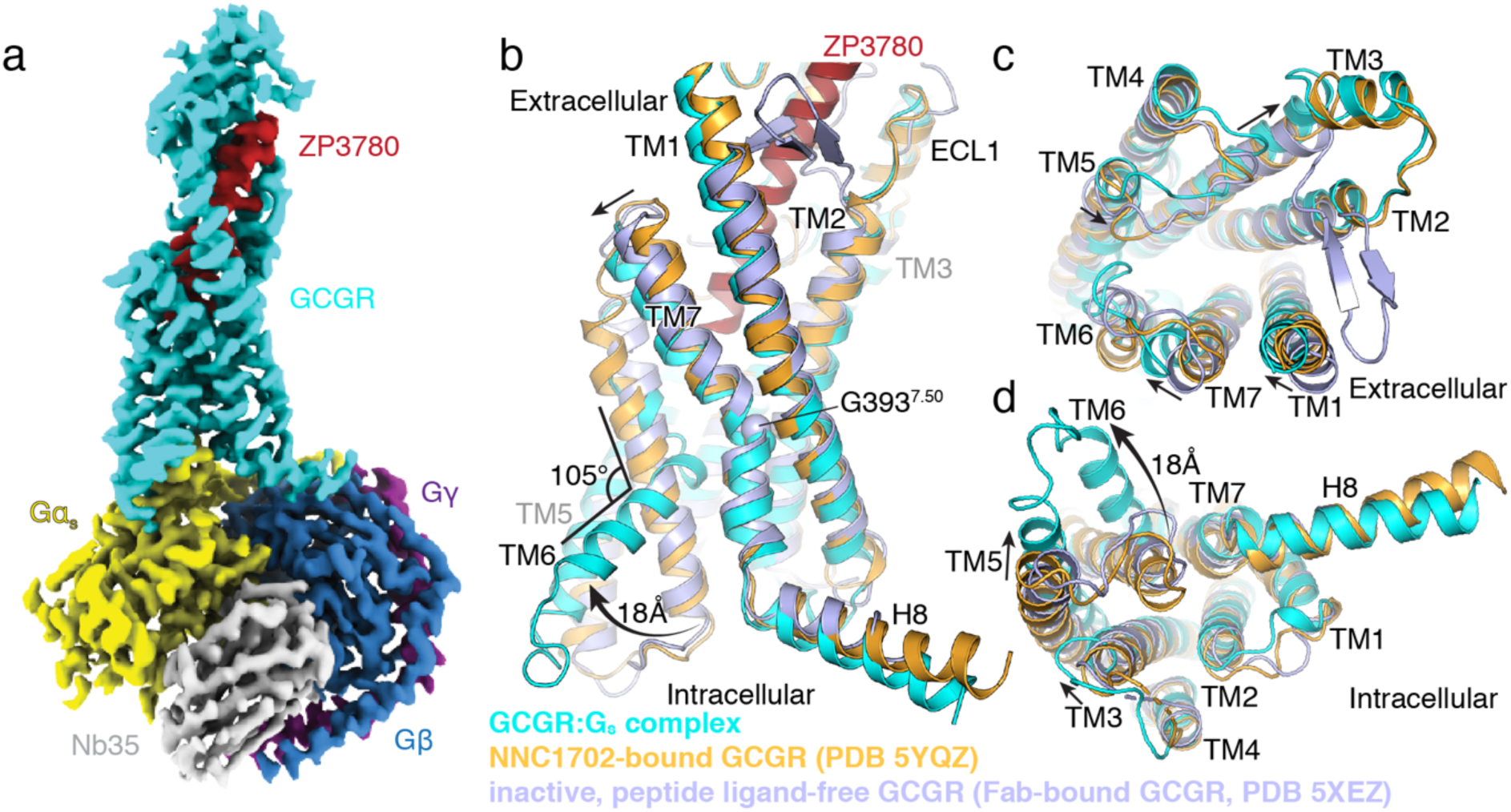
Cryo-EM structure of ZP3780-bound GCGR:G_s_ complex. a, Cryo-EM density map of the ZP3780-bound GCGR:G_s_ heterotrimeric complex colored by subunit. Cyan: GCGR, red: ZP3780, yellow: Gα_s_, blue: Gβ, purple: Gγ, grey: Nb35. b-d, Comparison of the inactive, peptide ligand-free state of GCGR (Fab-bound GCGR, blue, PDB 5XEZ), partial agonist NNC1702-bound GCGR (orange, PDB 5YQZ), and active, full-agonist ZP3780-bound state of GCGR (cyan). Significant structural change is observed in TM6, which moves outward by 18 Å and partially unwinds to form a kink with an angle of 105°. Additional changes are also observed in TMs 1, 3, 5, and 7.

The cryo-EM map shows well-defined density that allowed unambiguous modeling of the secondary structure and side chain orientations of all components of the GCGR-G_s_ complex, including the full agonist peptide ZP3780 (Suppl. Fig. 2, 3, 4a). In the structure, ZP3780 forms a continuous α-helix that is bound between the extracellular domain (ECD) and the transmembrane bundle of the receptor (Fig. 1a, Suppl. Fig. 4a, b). The peptide engages GCGR through extensive van der Waals and hydrophobic interactions with residues in the ECD and mostly polar contacts with residues in the transmembrane domain (Fig. 2a, Suppl. Fig. 4b-d). The overall peptide binding mode and the relative orientation of the ECD with respect of the transmembrane domain is very similar to the recently reported crystal structure of GCGR bound to the partial agonist peptide NNC1702 (Suppl. Fig. 4b)^10^. However, significant structural rearrangements are observed in the transmembrane region of the receptor (Fig. 1b-d). A comparison of the previously determined full-length GCGR structures in the inactive peptide-free (apo) state^9^ and bound to the partial agonist NNC1702^10^ with the fully active ZP3780-bound complex shows that peptide binding induces not only conformational changes of the ECD and the stalk region, but also at the extracellular ends of TMs 1, 2, 6, and 7 (Fig. 1b, c). These rearrangements in the transmembrane region widen the ligand binding site allowing the penetration of the peptide N-terminus deep into the receptor core. The GCGR:G_s_ complex structure also reveals structural changes that are related to full activation of the receptor. These include movement of the extracellular half of TM7 that bends further towards TM6 (6.5 Å C_α_-C_α_ distance between L377^7.34^) (receptor residues in superscript are defined using the class B numbering system)^20^, facilitated by the conserved G393^7.50^ in the center of the transmembrane domain (Fig. 1b, c). Furthermore, the entire TM3 shifts up by half a helix turn and its extracellular end moves away from the receptor core. On the intracellular side, TM5 moves towards TM6 by 6.5 Å, followed by the C-terminal end of TM3 (Fig. 1c,d). The most profound structural rearrangement between the NNC1702-bound structure of GCGR and the fully active receptor conformation in the G protein complex involves the formation of a sharp kink in the middle of TM6 (105.5°, measured between V368-G359-K344) (Fig. 1b). As reported for the previously determined active structures of other Class B GPCRs (GLP-1, CTR, CGRP, and PTH), the kink pivots the intracellular half of TM6 to move outwards by approximately 18Å, while the α-helical structure of the extracellular half partially unwinds ^12-16^. The conformational differences between the NNC1702 and the ZP3780-bound structures are most likely due to the distinct efficacy profiles of both ligands that are a result of sequence variations in the N-terminal part of the peptides, [as well as the presence of Gs]. In the native glucagon peptide, the N-terminal residues, histidine 1 (H1) and aspartate 9 (D9) have been shown to be critical positions for glucagon activity^11,21^. While those residues are present in ZP3780, NNC1702 exhibits a deletion of H1 and substitution of D9 by glutamate (D9E). Substitution of H1 or D9 to alanine in glucagon results in a marked reduction in ligand efficacy^22^. Therefore, a comparison between the interaction patterns of the N-termini of the peptides ZP3780 and NNC1702, and the transmembrane bundle (TMs 3, 5, 6, 7, and ECL3) of the receptor can provide a structural framework to explain the molecular basis of ligand efficacy for the glucagon receptor.

**Fig 2.**
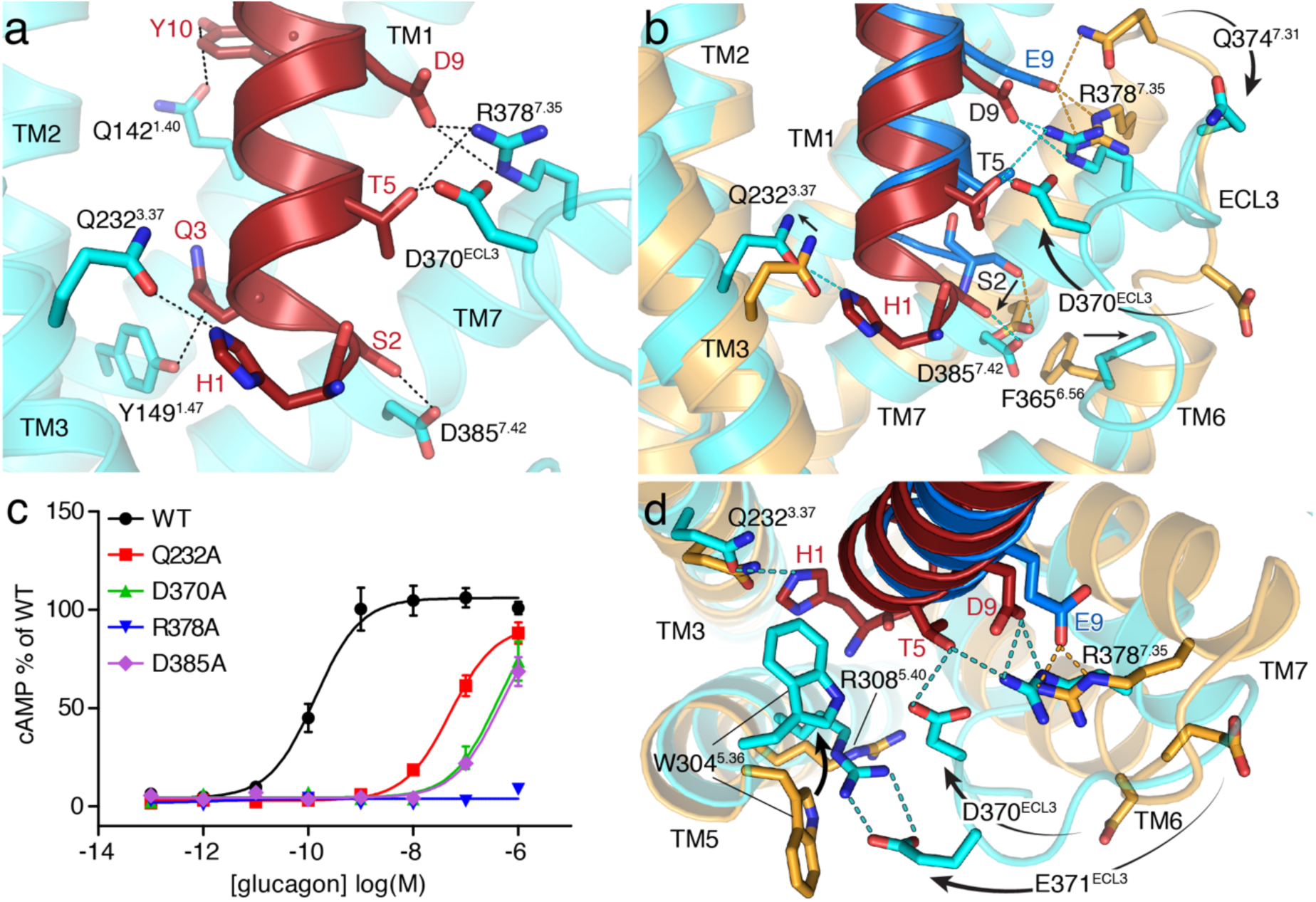
ZP3780 binding and activation at GCGR. a, The N-terminus of ZP3780 (red) is required for full ligand efficacy and penetrates deep in the receptor core to make hydrogen-bonds (H-bonds) (dotted lines) with residues in TMS 1, 3, 7 and ECL3 (cyan). b, The difference in receptor recognition by full-agonist ZP3780 (red) and partial-agonist NNC1702 (blue) that lacks H1 and has a D9E mutation is shown. NNC1702-bound GCGR is shown in orange (PDB 5YQZ) and the ZP3780-bound GCGR:G_s_ complex structure is shown in cyan. The polar interactions are shown as dotted lines colored based on the GCGR structures bound to the two ligands, respectively (ZP3780: cyan and NNC1702: orange). c, Mutation of Q232^3.37^, D370^ECL3^, R378^7.35^, and D385^7.42^d to alanine have a large effect on GCGR-mediated cAMP signaling. All mutants were expressed to similar levels as the WT receptor. d, Comparison of the interactions formed between the N-terminus of ZP3780 (red) and GCGR (cyan) and those formed by NNC1702 (blue) bound to GCGR (orange, PDB 5YQZ). The presence of H1 in ZP3780 ensures interaction with Q232^3.37^ and rearrangement of residues in TM5. The interaction of D9 stabilizes TM7 and ECL3 displacement, which might trigger GCGR activation. For c, data represent mean ± s.e.m. from at least three independent experiments, performed in triplicates.

In the full agonist ZP3780, the carboxyl group of D9 forms a salt bridge with residue R378^7.35^ on the N-terminus of TM7 that draws TM7 towards TM6 (Fig. 2a, b). This polar interaction is also present in previously determined GLP-1:G_s_ complex structures and in fact alanine mutation of R378^7.35^ completely inhibits GCGR-mediated cAMP signaling (Fig. 2c)^12,13,23^, thus highlighting the importance of this interaction for ligand-dependent activation within receptors of the glucagon family. The altered position of TM7 is accompanied by movement of ECL3 towards the receptor core wherein an hydrogen bond (H-bond) is formed between D370^ECL3^ and T5 of ZP3780 (Fig. 2b, d). This interaction is part of a polar network that includes R378^7.35^ and D9 and that stabilizes the active TM7 conformation together with the H-bond between S2 and D385^7.42^. Mutation of D370^ECL3^ and D385^7.42^ to alanine reduces ligand potency by at least 700-fold (Fig. 2c). The N-terminal residue H1 of ZP3780 forms a H-bond with residue Q232^3.37^ in TM3 on the other side of the ligand binding pocket. Formation of this interaction anchors the peptide deep into the ligand binding pocket and constrains its movement. Alanine substitution of Q232^3.37^ reduces the potency 350-fold, suggesting the importance of this interaction for ligand binding and/or receptor activation. Upon peptide binding, W304^5.36^ in TM5 flips towards the receptor core to stabilize the H1-Q232^3.37^ interaction by forming a “cap” that provides a further steric hindrance to peptide movement within the binding pocket (Fig. 2d). The functional importance of W304^5.36^ is shown by a mutation to glutamine that completely abolishes peptide binding^24^. Another consequence of the inward movement of W304^5.36^ seems to be to transmit the rotation through the helix to facilitate a large-scale translation of the cytoplasmic side of TM5 towards TM6 upon GCGR activation.

Comparison of the N-terminal positions of the full agonist ZP3780 and the partial agonist NNC1702 shows that the absence of H1 results in an upward shift of NNC1702 towards TM7 (Fig. 2b). In this position, the partial agonist would clash with TMs 1 and 7 in the fully activated receptor (Suppl. Fig. 5). In addition to this steric restriction of TM1 and TM7 movement, the D9E mutation allows the partial agonist to form a stronger contact with the extracellular end of TM7 (residues R378^7.35^ and Q374^7.31^), which stabilizes it more in an inactive state and prevents the inward movement of ECL3 (described above) (Fig. 2b, d). Furthermore, the missing N-terminal H1 and the lack of rearrangement in ECL3 would restrict the inward rotation of W304^5.36^, and as a consequence, the translational rotation of TM5 that is important for full activation of GCGR (Fig. 2d). Thus, comparison of the NNC1702 and ZP3780-bound GCGR structures provide insights into ligand-dependent conformational changes that might explain the molecular basis of ligand efficacy on the glucagon receptor. One hallmark of GPCR activation is the outward movement of TM6 that is accompanied in Class B GPCRs by the formation of a sharp kink in the middle of the transmembrane domain at the highly conserved PxxG motif^25^. In comparison to the NNC1702-bound receptor structure that shows an outward movement of the extracellular tip of TM6 induced by peptide binding, the ZP3780-bound GCGR:G_s_ complex reveals partial unwinding of helix 6 above the PxxG (P356^6.47^-L357^6.48^-L358^6.49^-G359^6.50^ in GCGR) motif to avoid spatial clashes with both the peptide and the repositioned TM7 (Fig. 3a, b; Suppl. Fig. 6). While this might initiate the destabilization of TM6 helicity, other conformational rearrangements around the PxxG motif seem to be ultimately required for kink formation. Ligand-induced movement of TM5 (discussed above) causes F322^5.54^ to relocate towards TM6 creating a clash between this bulky side chain and P356^6.47^ of the PxxG motif in the helical conformation of TM6, as seen in the NNC1702 bound structure (Fig. 3b)^10^. As a result, P356^6.47^ rotates away by approx. 90° causing the PxxG motif to shift inwards towards TMs 3 and 7 (Fig. 3a, b). Substitution of F322^5.54^ to alanine results in decreased potency of glucagon (800-fold), showing that the clash of the PxxG motif with TM5 is important for receptor activation (Fig. 3c). Furthermore, a slight rotation of TM7 away from TM6, caused by the ligand-induced changes described above, helps to accommodate the displaced PxxG motif in the active state of the receptor by releasing the steric hindrance between the newly positioned P356^6.47^ and L395^7.52^ in TM7 (Fig. 3b). Another obstacle for kink formation is found in TM3, where the side chain of L242^3.47^ would prevent inward movement of the PxxG motif. The observed upward shift of TM3 leads to the removal of this restriction by shifting the leucine side chain away from the PxxG motif (Fig. 3b). Individual replacement of the two leucines (L242^3.47^ and L395^7.52^) in TMs 3 and 7 to the smaller alanine side chain leads to a nearly 20-fold decrease in potency and a significant decrease in the maximal glucagon response (Fig. 3c, Suppl. Table 6). This highlights the importance of the conformational rearrangements necessary for the inward rotation of the PxxG motif upon receptor activation. The discussed repositioning of the PxxG motif is further stabilized by H-bonds with TM5 (G359^6.50^-N318^5.50^) and TM7 (P356^6.47^-Q392^7.49^) (Fig. 3b).

**Fig 3.**
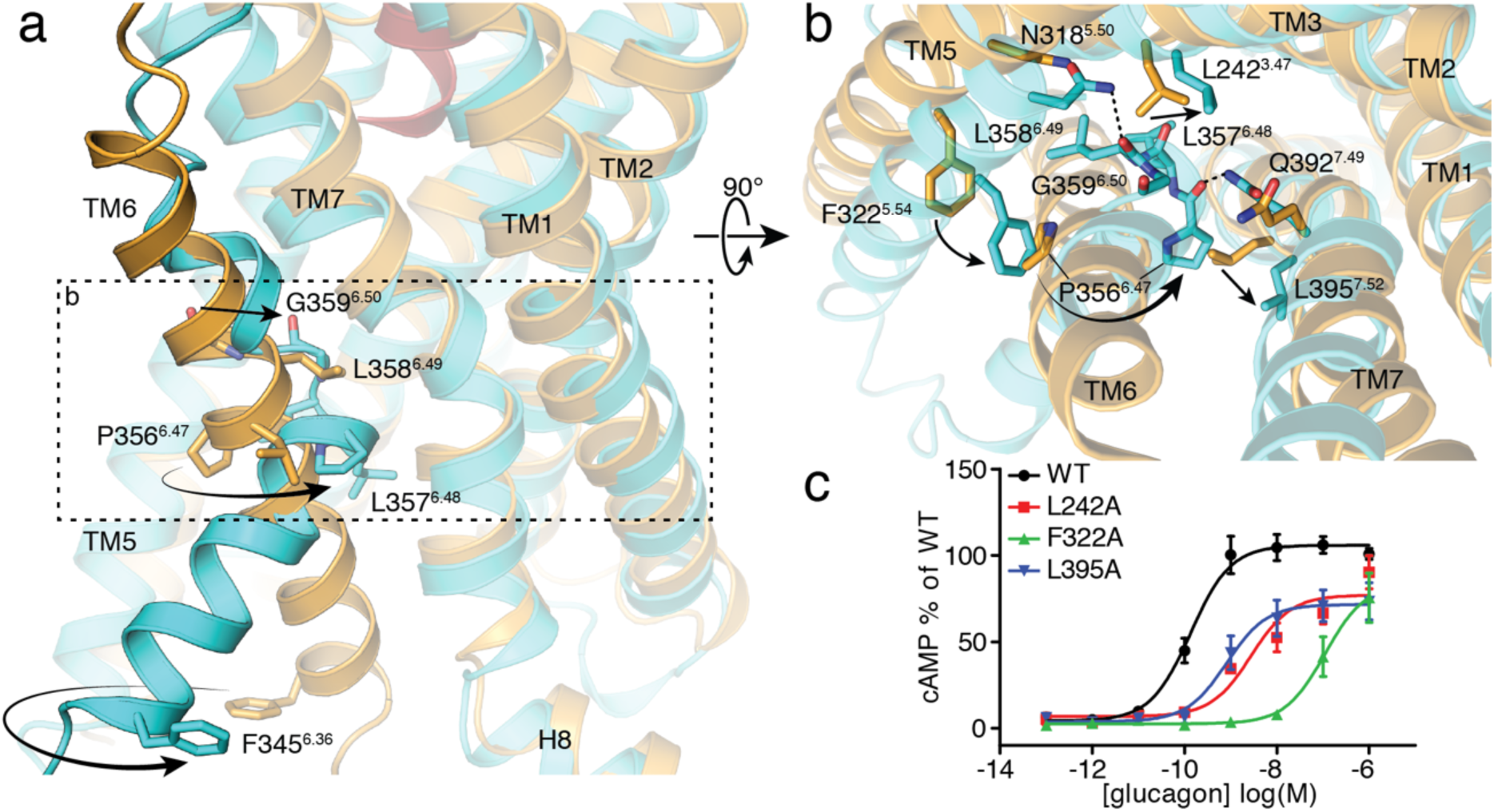
Structural changes in the PxxG motif of GCGR induced by full-agonist ZP3780 and G_s_ coupling. a, The rearrangement of TMs 3, 5 and 7 allows TM6 outward movement and ‘kink’ formation at the conserved PxxG motif (P356^6.47^-L357^6.48^-L358^6.49^-G359^6.50^). The ZP3780-bound GCGR:G_s_ complex (cyan) is overlaid with that of the NNC1702-bound GCGR (orange) to highlight structural changes that results in the TM6 outward movement in the GCGR:G_s_ complex structure. b, Reorientation of residues in TMs 3, 5, and 7 around the PxxG motif that facilitate kink formation in TM6. c, Alanine substitution of residues around the PxxG motif significantly reduce GCGR-mediated cAMP signaling. All mutants were expressed to similar levels as the WT receptor. For c, data represent mean ± s.e.m. from at least three independent experiments, performed in triplicates.

Below the PxxG motif, the 90° rotation of P356^6.47^ perpetuates towards the N-terminus of TM6 that moves away from TM5 maintaining the 90° rotation and is displaced from the receptor core by 18Å to engage the G protein (Fig. 1b, Fig. 4a). The conformational alterations in the transmembrane bundle of the receptor open up an intracellular cavity that allows engagement of the heterotrimeric G protein, G_s_. The overall structure of the ZP3780-GCGR:G_s_ complex displays high similarity in the receptor-G protein interactions compared to other reported structures of Class B GPCRs-G_s_ complexes (GLP-1, Calcitonin, CGRP, PTH), suggesting a common mechanism for G protein engagement and activation (Suppl. Fig. 7)^13-16,26,27^. In the nucleotide-free GCGR:G_s_ complex, the position of the α5 helix of the G protein requires outward movement of TMs 5 and 6 due to steric hindrance (Fig. 4b). However, superimposition of the inactive GDP-bound G_s_ heterotrimer^28^ onto the nucleotide-free G_s_ in the complex suggests that the bent α5 helix of the GDP-bound G protein (α5^GDP^) together with the known inherent flexibility of its very C-terminal end^29,30^, might allow the initial engagement of the GDP-bound G protein without outward displacement of TM6 (Fig. 4b). In this orientation of α5^GDP^, TM5 would need to relocate in order to prevent clashes, but would still allow the establishment of the polar interactions between α5 (e.g. R385) and TM5 as seen in the nucleotide-free state (Fig. 4). These interactions might facilitate the straightening of α5 upon nucleotide release accompanied by outward movement of the intracellular tip of TM6 leading to the sharp kink as seen in GCGR and other Class B GPCR complexes. As a result, α5 inserts more deeply into the receptor core to form extensive hydrophilic interactions with residues in TMs 5 and 6 and helix 8 (H8) and hydrophobic contacts with TM3 (Fig. 4c). Additional stabilization of the outward conformation of TM6 is provided by rearrangements in the highly conserved HETx motif just below the kink region that has been shown to be important for receptor activation (Suppl. Fig. 8)^25,31^. Removal of the hydroxyl of Y400^7.57^ in the HETx motif leads to a 40 percent decrease in maximal cAMP signaling by glucagon as compared to the wild type (Suppl. Fig. 8b), suggesting the critical role of this polar network for receptor activation.

**Fig 4.**
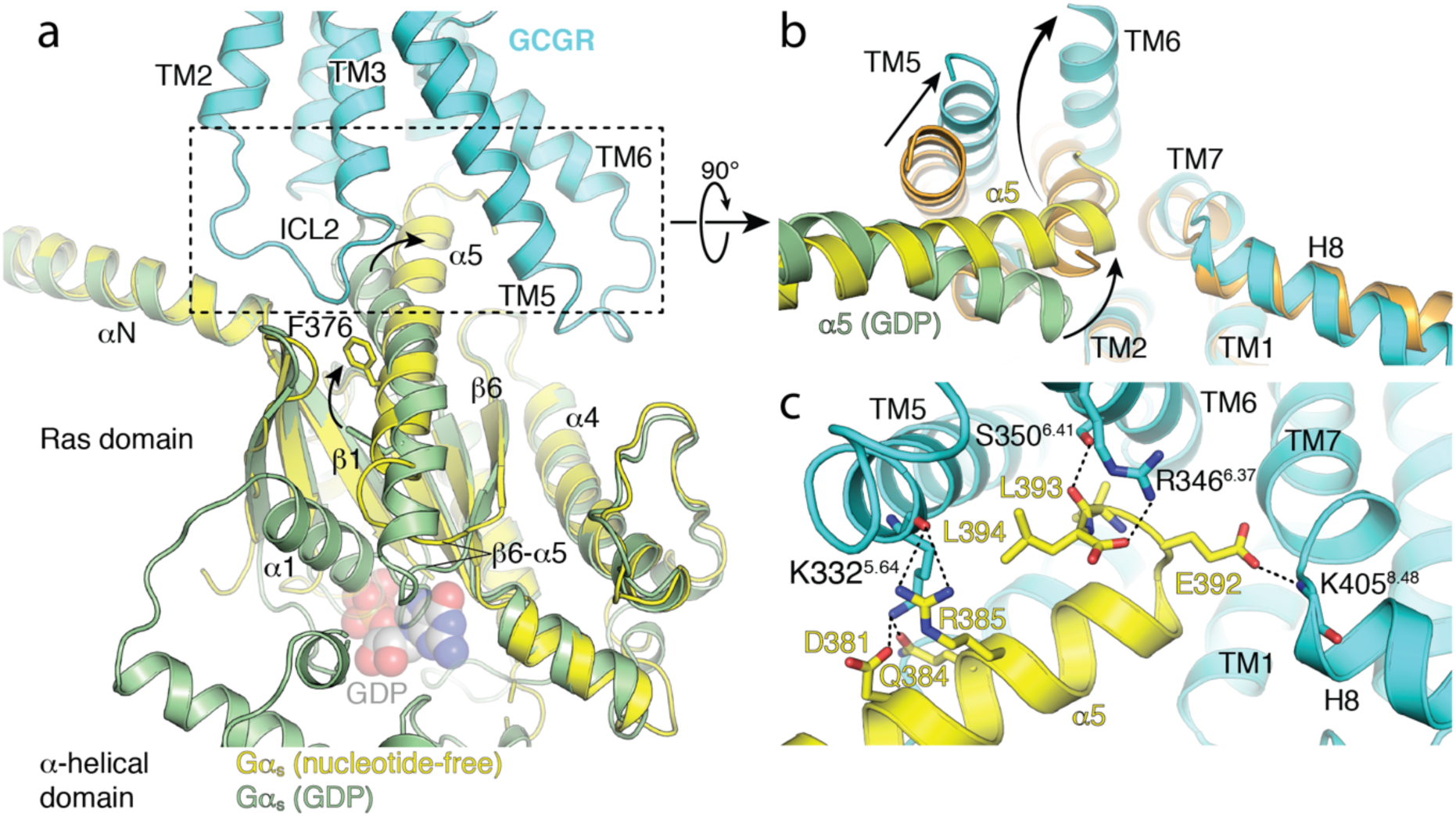
Structural changes in G_s_ on GCGR binding. Comparison of GDP-bound Gα_s_ (green, PDB 6EG8)^28^ and nucleotide-free Gα_s_ (yellow) in complex with GCGR (cyan) showing differences in the α1 helix, α5 helix, and α5-β6 loop as well as in the α-helical domain upon binding to the receptor. b, The initial α5 engagement of GDP-bound Gα_s_ (green) to GCGR with an inwardly positioned TM6 (orange, PDB 5YQZ) appears to be possible without the outward movement of TM6. The final conformation of the nucleotide-free GCGR:G_s_ complex (GCGR: cyan, Gα_s_: yellow) is achieved when α5 displaces TM6 outward (cyan), resulting in kink formation of the transmembrane domain. c, The α5 helix (yellow) of Gα_s_ in the nucleotide-free GCGR:G_s_ complex engages the cytoplasmic core of GCGR (cyan) to form extensive polar interactions (dotted line) with TMs 5, 6 and helix 8 (H8).

Another important interface between the receptor and the G protein, that has been shown to be important for G protein-signaling of GCGR, is mediated by ICL2 and 3^32^. While ICL3 interacts with residues in the α4/β6 region of the Ras-like domain, ICL2 points into a mostly hydrophobic pocket lined by residues in the αN/β1 hinge and α5 of Gα_s_ (Suppl. Fig. 9a,b). Furthermore, two arginine residues (R413^8.56^ and R417^8.60^) in H8 are shown to form H-bonds with backbone atoms of residues A309 and H311 in the WD7 blade of Gβ subunit (Suppl. Fig. 9c). This H8-Gβ interaction seems to be more conserved among Class B receptor:G protein complexes and has been shown to mediate the efficiency of G protein-dependent activation of the adenylyl cyclase^14^.

## Conclusions

To gain insights into the mechanism of ligand-induced activation of GCGR we designed a glucagon analogue with better aqueous solubility, ZP3780, and determined the structure of a GCGR:G_s_ signaling complex. The peptide residues H1 and D9, have been shown to be indispensable for ligand potency and efficacy^11,21^. This study provides a structural rationale to explain the wealth of existing structure-activity relationship data for glucagon peptides that has accumulated since the discovery of glucagon by Kimball and Murlin almost 100 years ago^33^. This study will allow the exploration of new avenues for the development of innovative therapeutics to modulate GCGR signaling and its impact on the pathophysiology of diabetes.

## Methods

### Solubility assay

Stock solutions of 2.0 mg/ml native glucagon and ZP3780 were prepared by dissolving the compounds in milliQ water adjusted to pH 2.5 by HCl. The concentration of compound was determined by absorbance measurement at 280 nm (Nanodrop 2000c, Thermo Scientific) using a calculated extinction coefficient of 8480 M^-1^cm^-1^. Aliquots of 50 μl buffer (Suppl. Table 1) were transferred to a clear bottom, UV compatible 96 wells plate and mixed with 50 μl aliquots of the peptide stock. The final peptide concentrations were 1 and 0.5 mg/ml and the final buffer concentration 50 mM. ZP3780 was also determined at 5 mg/mL using a stock solution of 10 mg/mL, The samples were incubated at room temperature for 15 minutes and the UV absorbance at 325 nm was measured by an Elisa reader (Spectramax 190). The samples were considered fully dissolved if the absorbance at 325 nm was below 0.02 AU, which corresponds to 5 to 6 times the standard deviation of 8 buffer samples.

### Aggregation assay

Samples were prepared according to the solubility assay, with the addition of a final concentration of 40 µM ThT. Samples were loaded in a 96-well black fluorescence plate (clear bottom) in triplicates of 150 µL in each well at ambient temperature and subsequently the plate was covered with UV compatible sealing tape to avoid evaporation. The plates were placed in a Spectramax Gemini XS fluorescence plate reader. The plate reader was programmed to read fluorescence from the top of the plate at 355 nm (excitation 295 nm) and at 485 nm (excitation 450 nm) at fixed intervals of 10 min preceded by 300 s of automixing for 96 h at 40° C.

### ^125^I-glucagon competition binding assay

Membranes containing the human GCGR (hGCGR) were prepared as described34 from a HEK293 cell line stably over-expressing the hGCGR. In brief, for generation of the over-expressing cell line, cDNA encoding the hGCGR (P47871) was amplified by PCR using primers encoding terminal restriction sites for subcloning. The 5’-end primer additionally contained a near Kozak consensus sequence. The fidelity of the DNA encoding the hGCGR was confirmed by DNA sequencing. The PCR product encoding the hGCGR was subcloned into a mammalian expression vector containing a neomycin (G418) resistance marker. The mammalian expression construct was transfected into HEK293 cells by a standard liposome transfection method. 48 hours post-transfection, cells were seeded for limited dilution cloning and selected with 0.5 mg/ml G418 in the culture medium. After 2 weeks, surviving colonies of hGCGR-expressing cells were picked, propagated and tested for cAMP accumulation. One hGCGR-expressing clone was selected and used for membrane preparation.

For the competition binding experiments 0.06 nM [^125^I]-glucagon (PerkinElmer, NEX207) was added together with 10 µg hGCGR containing membranes, 0.5 mg SPA-beads (PerkinElmer, RPNQ0001) and varying concentrations of native glucagon or ZP3780. After sharing at 600 rpm for 2 h at room temperature the plate transferred to a MicroBeta2 scintillation counter (PerkinElmer) left for 4 h to allow the SPA-beads to settle before quantifying ^125^I radioactivity. To determine the respective K_i_-values, data were fitted by nonlinear regression to a one-site competition model using GraphPad Prism.

### Functional cAMP accumulation assay

The in vitro potencies and efficacies of glucagon and ZP3780 were assessed in a cAMP accumulation assay using HEK293 cells that were transiently transfected with the hGCGR. In brief, HEK293 cells were brought into suspension (0.25×10^6^ cells/mL) by trypsination. For transfection of 1 mL of cells in suspension, a total DNA amount of 1 µg, consisting of 0.2 µg hGCGR (accession no P47871) expression vector DNA and 0.8 µg pcDNA3.1(zeo), in 25 µl OptiMEM was mixed with 3 µl FuGene6 in 57 µL OptiMEM and incubated for 20 min. Hereafter the 85 µL DNA/FuGene6 mix was added to 1 mL cell suspension, mixed and seeded in a clear poly-D-lysine coated 96-well in a volume of 100 µL/well (final cell density of 2.5×10^5^ cells/well). The amounts were scaled up to the number of data points needed to generate concentration response curves for glucagon and ZP3780. Twenty-four hours after transfection the assay was performed by washing wells once in 100 µL HBSS buffer (HBSS Gibco 14025, 20 mM HEPES pH 7.5, 1 mM CaCl_2_, 1 mM MgCl_2_ and 0.1% BSA). After the wash, 100 µL/well of HBSS buffer supplemented with IBMX (100 µM final concentration) were added, quickly followed by addition of 50 µL of HBSS buffer containing compound dilutions of glucagon and ZP3780. After incubation at 37°C for 15 min, the reactions were stopped by addition of 160 µL lysis buffer. The accumulated cAMP levels were quantified using the CisBIO cAMP dymanic HTRF kit (Cisbio, 62AM4PEC) according to the manufacturer’s instructions and measured on an EnVision plate reader (PerkinElmer). Data were fitted by non-linear regression using GraphPad Prism.

Functional testing of the hGCGR mutations was, in general, conducted as described above. The constructs used for the mutational studies all contained a N-terminal FLAG-tag preceded by a HA-signal peptide and were inserted into the pcDNA3.1(neo) expression vector. The following DNA amounts were transfected to ensure similar surface expression to WT hGCGR: WT GCGR was transfected with 0.2 µg/mL cell suspension, Q232A and D385A GCGR with 0.05 µg DNA/mL cell suspension, and finally L242A, F322A, D370A, R378A, L395A, and Y400F GCGR were transfected with 0.035 µg/mL cell suspension. Again, in all cases, the total DNA amount was adjusted to 1 µg/mL cell suspension by addition of pcDNA3.1(zeo). Twenty-four hours after transfection the assay was performed by washing wells once in 100 µL DPBS + 1 mM CaCl_2_ (DPBS Gibco 14190). After the wash, 50 µL/well HBSS buffer (HBSS Gibco 14025, 20 mM HEPES pH 7.5, 1 mM CaCl_2_, 1 mM MgCl_2_ and 0.1% BSA) was added and the plates were incubated for 30 min at 37°C. Hereafter, 50 µl/well glucagon dilutions were added, which were prepared in HBSS buffer supplemented with IBMX to a final concentration of 100 µM. After incubation at 37°C for 15 min, the reactions were stopped and the accumulated cAMP levels were quantified as described above.

### Enzyme-linked immunosorbent assay

Surface expression levels of WT and mutated hGCGR variants were tracked using a direct enzyme-linked immunosorbent assay (ELISA) against the N-terminal FLAG tag. In brief, cells were transfected and seeded in a white Poly-D-Lysine-coated 96-well plate as described for the functional cAMP accumulation assay. Twenty-four hours after transfection, cells were washed with 100 µl/well DPBS + 1 mM CaCl_2_ and fixed with 50 µl/well 4% paraformaldehyde solution for 5 minutes at room temperature. The wells were washed twice with 100 µl DPBS + 1 mM CaCl_2_ and blocked with 100 µl/well blocking solution (3% drymilk, 1 mM CaCl_2_, 50 mM Tris-HCl, pH 7.5) for 30 minutes at room temperature. Hereafter 75 µl/well HRP-conjugated anti-FLAG antibody (Sigma Aldrich, A8592), diluted 1:2000 in blocking solution, was added and the plate was incubated for 1 hour at room temperature. The plate was washed four times with 100 µl/well blocking solution followed by four washes with 100 µl/well DPBS + 1 mM CaCl_2_. The relative amount of present HRP was detected by adding 60 µL/well DPBS + 1 mM CaCl_2_ and 20 µl/well HRP substrate (Bio-Rad, 170-5060). The plate was incubated for 10 minutes at room temperature before recording of luminescence on an EnVision plate reader (Perkin Elmer).

### Purification of GCGR

Human GCGR (Q27-F477) with N-terminal FLAG and C-terminal octahistidine tag was expressed in *Spodoptera frugiperda Sf9* insect cells using the baculovirus method (Expression Systems) in the presence of the L-168,049 (Tocris Bioscience). GCGR was extracted with 1% lauryl maltose neopentyl glycol (L-MNG, Anatrace) and purified by nickel-chelating sepharose chromatography in the presence of the ligand NNC0640. The eluate from the nickel resin was applied to the M1 anti-FLAG immunoaffinity resin and washed with progressively decreasing concentration of NNC0640. The receptor was eluted in a buffer consisting of 20 mM HEPES pH 7.5, 150 mM NaCl, 0.05% L-MNG, 0.005% cholesterol hemisuccinate (CHS), FLAG peptide and 5 mM EDTA. The final step of purification was size exclusion chromatography on Superdex 200 10/300 gel filtration column (GE healthcare) in 20 mM HEPES pH 7.5, 150 mM NaCl, 0.02% MNG and 0.002% CHS. Finally, apo-GCGR was concentrated to ∼400 µM and stored at −80° C.

### Expression and purification of heterotrimeric G_s_

Heterotrimeric G_s_ was expressed and purified as previously described^35^. Briefly, heterotrimeric G_s_ was expressed in High Five™ (*Trichuplusia ni*) insect cells using baculoviruses generated by the BestBac method (Expression Systems). Two separate baculoviruses were used, one encoding the human Gα_s_ short splice variant and the other encoding both the Gβ_1_ and Gγ_2_ subunits, with an histidine tag and HRV 3C protease site inserted at the amino terminus of the β-subunit. High Five™ cells were infected with the baculoviruses followed by an incubation of 48 h at 27 °C. Cells were harvested by centrifugation and lysed in a buffer comprised of 10 mM Tris, pH 7.5, 100 μM magnesium chloride (MgCl_2_), 5 mM β-mercaptoethanol (βME), 50 μM GDP and protease inhibitors. The membrane fraction was collected by centrifugation and solubilized with a buffer comprised of 20 mM HEPES, pH 7.5, 100 mM sodium chloride (NaCl), 1% sodium cholate, 0.05% DDM, 5 mM MgCl_2_, 5 mM βME, 5 mM imidazole, 50 μM GDP and protease inhibitors. The solubilization reaction was incubated for 45 min at 4 °C after homogenization with a Dounce homogenizer. After centrifugation, the soluble fraction was loaded onto Ni-chelated sepharose followed by a gradual detergent exchange into 0.1 % DDM. The protein was eluted in buffer supplemented with 200 mM imidazole and dialyzed overnight in 20 mM HEPES, pH 7.5, 100 mM NaCl, 0.1% DDM, 1 mM MgCl_2_, 5 mM βME and 50 μM GDP together with HRV 3C protease to cleave off the amino-terminal 6xHis tag. The cleaved 6xHis tag, uncleaved fractions and 3C protease were removed by Ni-chelated sepharose and the G protein was dephosphorylated by lambda protein phosphatase (NEB), calf intestinal phosphatase (NEB), and antarctic phosphatase (NEB). Lipidated G_s_ heterotrimer was isolated using a MonoQ 10/100 GL column (GE Healthcare). After binding of the protein to the column in buffer A (20 mM HEPES, pH 7.5, 50 mM NaCl, 1 mM MgCl_2_, 0.02% DDM, 100 μM TCEP and 20 μM GDP), the column was washed with buffer A and the G protein heterotrimer was eluted with a linear gradient of 0–30% buffer B (buffer A with 1 M NaCl). The main peak containing isoprenylated G protein heterotrimer was collected and the protein was dialyzed into 20 mM HEPES, pH 7.5, 100 mM NaCl, 0.02% DDM, 100 μM TCEP and 50 μM GDP. After concentrating the protein to 250 μM, 20% glycerol was added and the protein was flash frozen in liquid nitrogen and stored at –80 °C until use.

### Purification of Nb35

Nb35 was expressed in *E. coli* BL21 cells. After lysis, it was purified on a nickel affinity chromatography and finally subjected to size exclusion chromatography on a Superdex 200 10/300 gel filtration column (GE healthcare) in 20 mM HEPES pH 7.5, 150 mM NaCl. It was flash frozen and stored at −80 °C.

### Formation and purification of the GCGR-G_s_-Nb35 complex

Purified G_s_β_1_γ_2_ in DDM was incubated with 1% MNG for one hour on ice and simultaneously, GCGR was incubated with 5-fold molar excess ZP3780 (500 µM) at room temperature. A 1.5-fold molar excess of detergent-exchanged Gα_s_β_1_γ_2_ was incubated with ZP3780-bound GCGR at room temperature for two hours, after which a 2-fold molar excess (in terms of G_s_β_1_γ_2_) of Nb35 was added and incubated on ice for 1.5 hours. To stabilize the nucleotide-free complex, Apyrase (1 unit, NEB) was added and incubated overnight at 4° C. A 4-fold volume of 20 mM HEPES pH 7.5, 100 mM NaCl, 0.8% L-MNG/0.08% CHS, 0.27% GDN/0.027% CHS, 1 mM MgCl_2_, 5 µM ZP3780, and 2 mM CaCl_2_ was added to the complexing reaction and complex was purified by M1 anti-FLAG affinity chromatography. The complex was eluted in 20mM HEPES pH 7.5, 100mM NaCl, 0.01% MNG/0.001% CHS, 0.0033% GDN/0.00033% CHS, 5 µM ZP3780, 5 mM EDTA, and FLAG peptide. The eluted complex was supplemented with 100 µM TCEP and subjected to size exclusion chromatography on a Superdex 200 10/300 Increase column in 20mM HEPES pH 7.5, 100mM NaCl, 5 µM ZP3780, 0.00075% MNG and 0.00025% GDN. Peak fractions were concentrated to ∼16 mg/mL for electron microscopy studies.

### Cryo-EM data acquisition and data processing

An aliquot of 3.5 μL GCGR-G_s_-Nb35 complex was applied to glow-discharged 200 mesh grids (Quantifoil R1.2/1.3), at a concentration of 17 mg/ml and subsequently vitrified using a Vitrobot Mark IV (Thermo Fischer Scientific) at 100 % humidity and 4° C. Cryo-EM images were collected on a Titan Krios operated at 300 kV at a nominal magnification of 130,000× with a Gatan GIF Quantum LS Imaging energy filter using a Gatan K2 Summit direct electron camera in counted mode, corresponding to a pixel size of 1.06 Å. A total number of 3724 movie stacks were obtained with a dose rate of 7 e^-^/pix/s and total exposure time of 8 s with 0.2 s per frame, resulting in a total dose of 50 electron per Å^2^. The defocus range was set to 1.2-2.2 μm. Dose-fractionated image stacks were subjected to beam-induced motion correction using MotionCor2^36^. Contrast transfer function parameters for each micrograph were determined by Gctf v1.06^37^. Data processing was performed in RELION3.0^38^. A total number of 2,039,910 particles were selected from a template-based auto-picking. A subset of 296,516 particles were selected after one round of 2D and 3D classification. Particles projections from micrographs with signal better than 3.5Å were subjected to Bayesian polishing, CTF refinement and 3D reconstruction. The final subset of 266,267 particles was imported to cisTEM for 3D local refinement, resulting in a reconstruction with global resolution of 3.1Å at FSC 0.143^39^. Local resolution was determined using the Bsoft package at cutoff FSC of 0.5^40^.

### Model building and refinement

The initial template of GCGR was derived from the crystal structure of GCGR (PDB 5YQZ)^10^. The receptor coordinates (PDB 5VAI) ^27^ were used as initial models for the G_s_ and Nb35. Models were docked into the EM density map using *UCSF Chimera*^41^, followed by iterative manual building in *Coot*^42^. The final model was subjected to global refinement and minimization in real space using *phenix.real_space_refine* in *Phenix*^43^. *Molprobity* was used to evaluate model geometry^44^. FSC curves were calculated between the resulting model and the half map used for refinement as well as between the resulting model and the other half map for cross-validation.

### Figure preparation

Figures were created using the *PyMOL* Molecular Graphics System, Version 2.20 Schrödinger, *LLC* (http://pymol.org/), and the UCSF *Chimera X* package^45^. In vitro graphs were created using *GraphPad Prism*.

## Acknowledgments

D.H. was supported by the German Academic Exchange Service (DAAD). K.K. was supported by the American Diabetes Association (ADA) Postdoctoral Fellowship. The work is supported by NIH grant R01GM083118. B.K.K. is a Chan Zuckerberg Biohub investigator. We thank Evan O’Brien for critical reading of the manuscript.

