## Supplemental data for "Structural insights into ligand efficacy and activation of the glucagon receptor"

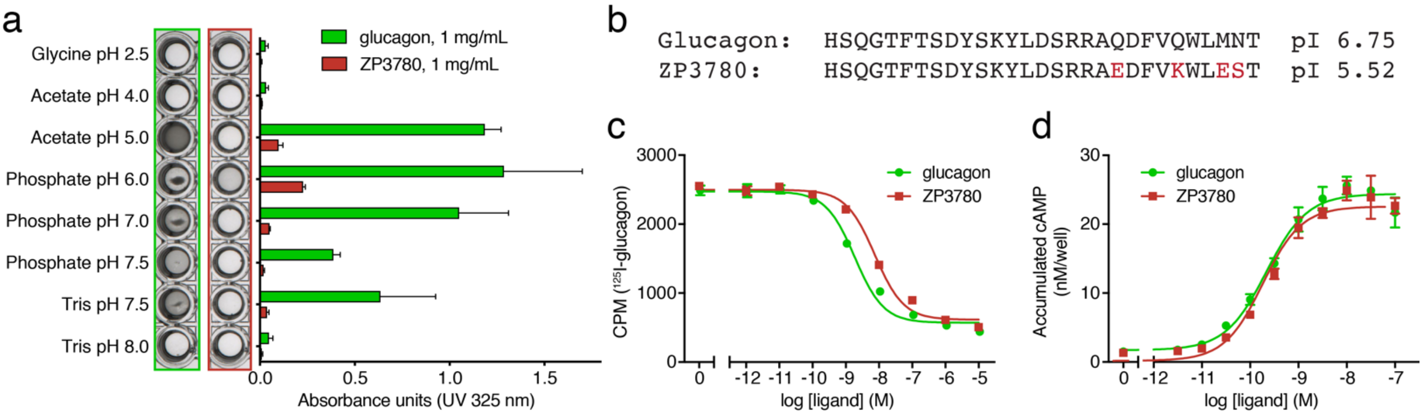


**Supplemental Fig. 1:** ***In vitro*** **properties of ZP3780 compared to glucagon.** a, The solubility of ZP3780 (red) is improved compared to glucagon (green) in a large range of buffers and pH, also around pH 7.5 as observed and measured by UV 325 nm absorbance. b, Sequence of glucagon and ZP3780. c, ZP3780 (red) displaces $^{125}$I-glucagon to non-specific levels as observed for glucagon (green) in competition binding studies with GCGR overexpressed in HEK293 cell membranes. d, ZP3780 (red) has a similar potency and Emax as glucagon (green) in a functional cAMP accumulation assay in HEK293 cells transiently expressing the GCGR. The affinity of ZP3780 (c) is marginally lower than glucagon (also refer to Suppl. Tables 3, 4). Data represent mean ± s.e.m. from three (a) or four (c and d) independent experiments.

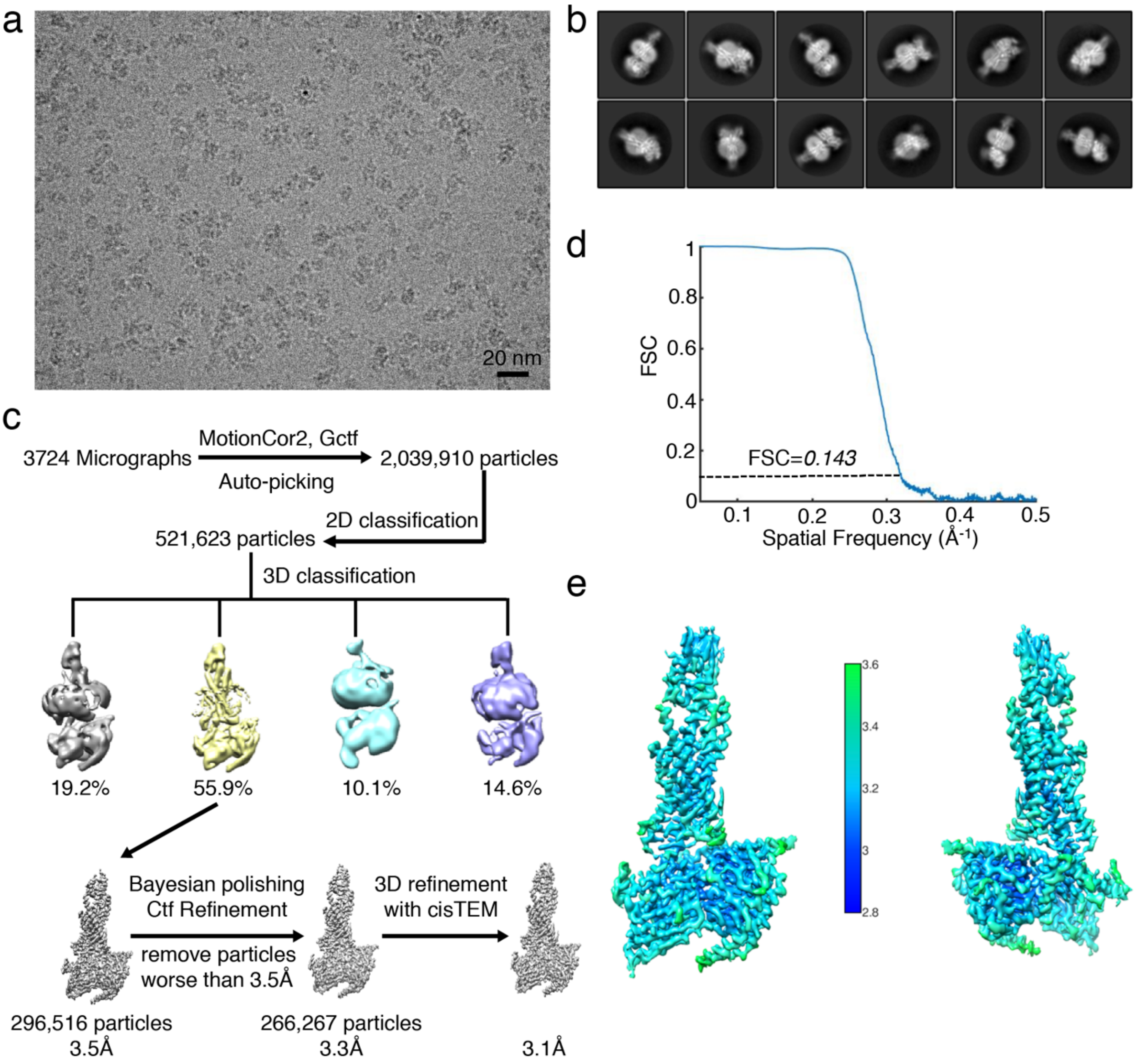


**Supplemental Fig. 2: Cryo-EM data processing.** a, Representative cryo-EM image of the GCGR:$G_s$ complex. Scale bar: 20 nm. b, Representative reference-free 2D cryo-EM averages of GCGR:$G_s$. c, Cryo-EM data processing flow chart of GCGR, including particle selection, classifications and density map reconstruction. d, 'Gold standard' FSC curves from RELION[1] indicate that the map for the GCGR:$G_s$ complex reaches a nominal resolution of 3.1Å at FSC = 0.143. e, Three-dimensional density maps of the GCGR:$G_s$ complex, colored by local resolution.

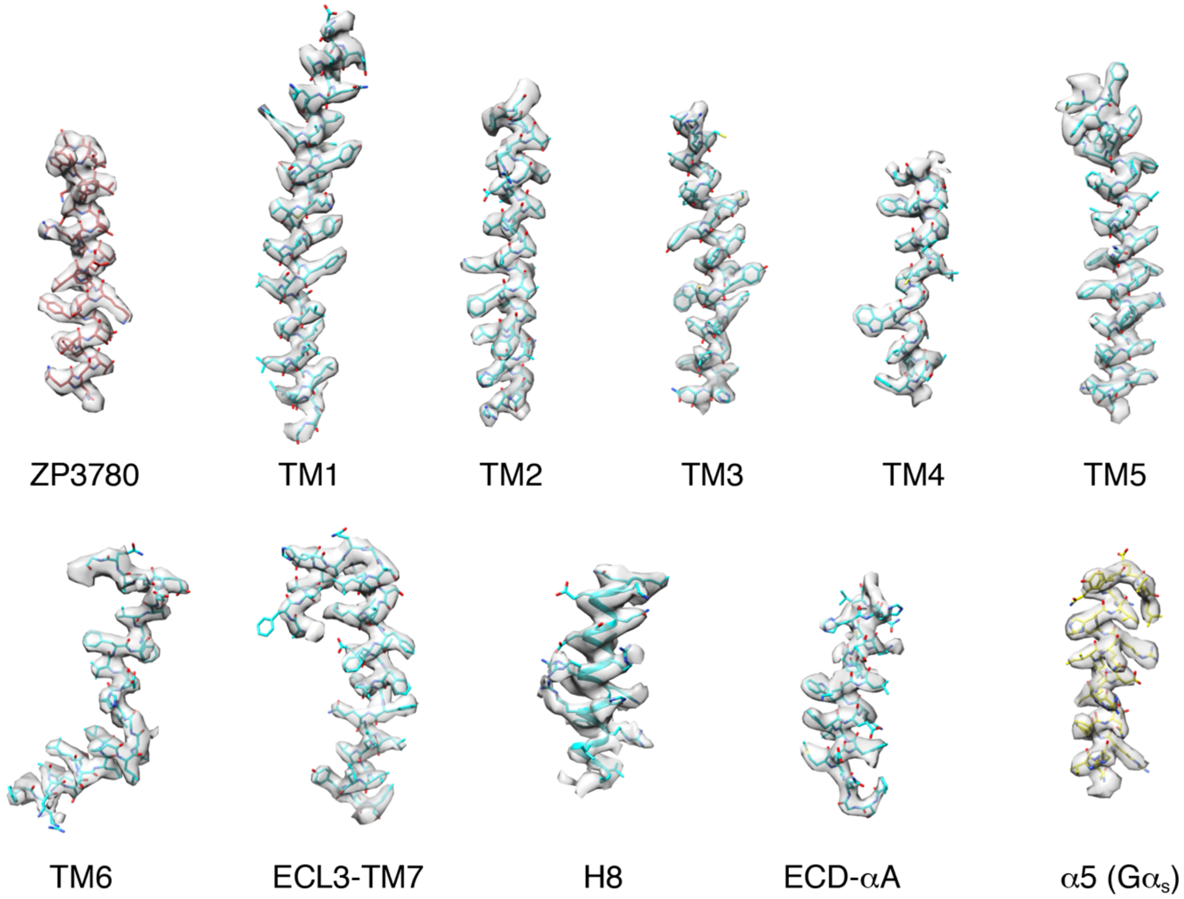


**Supplemental Fig. 3: Map/model quality and local resolution.** Cryo-EM density map and model are shown for ZP3780, all seven transmembrane α-helices, ICL3, and αA (ECD) of GCGR and α5 of Gα$_s$.

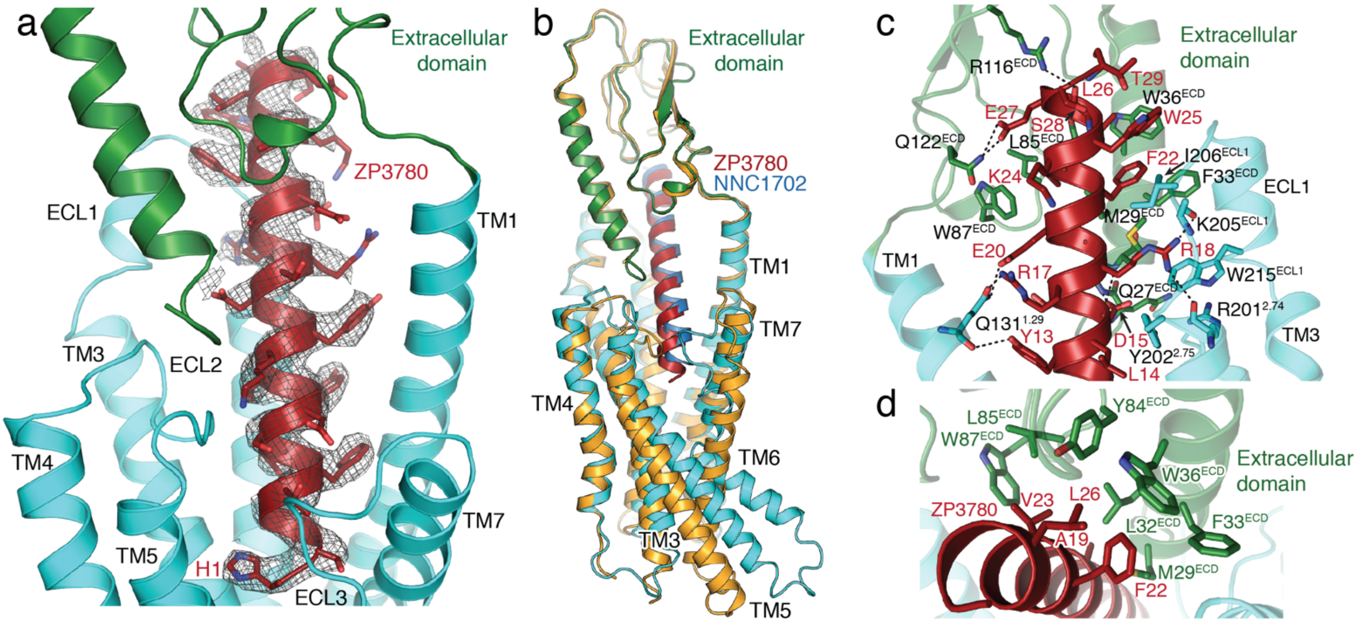


**Supplemental Fig. 4: Interactions between the full agonist ZP3780 and GCGR.** a, The cryoEM density (displayed as a mesh) of the bound ZP3780 (red) in the peptide binding pocket of GCGR (TMs shown in cyan and the extracellular domain (ECD) shown in green). b, Comparison of peptide-binding mode of full-agonist ZP3780 (red)-bound GCGR (cyan) and partial agonist-bound NNC1702 (blue)-bound GCGR (orange) (PDB 5YQZ)[2]. c, The C-terminus of ZP3780 (red) engages residues in polar (dotted lines) and hydrophobic (residues shown) interactions with the ECD (green), TMs 1, 2, 3, and ECL1 (cyan). d, Hydrophobic interaction between residues in ZP3780 (red) and the ECD of GCGR (green).

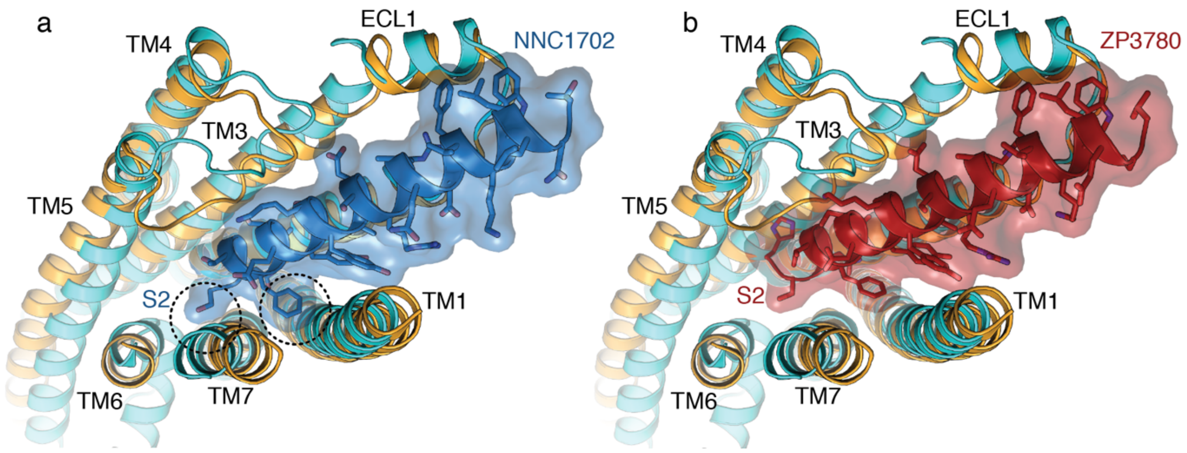


**Supplemental Fig. 5: Comparison of the binding mode of full- and partial agonists to GCGR.** a, The binding pose of the peptide in the partial agonist NNC1702 (blue)-bound GCGR structure (orange, PDB 5YQZ)[2] would cause steric clashes with TMs 1 and 7 in the active state of GCGR (highlighted with dashed circles) in the GCGR:$G_s$ complex structure (cyan). b, The full agonist ZP3780 in the GCGR:$G_s$ complex structure (cyan) binds closer to TM3 resulting in a relief of the clashes with TMs 1 and 7 seen in (a) with the partial agonist NNC1702.

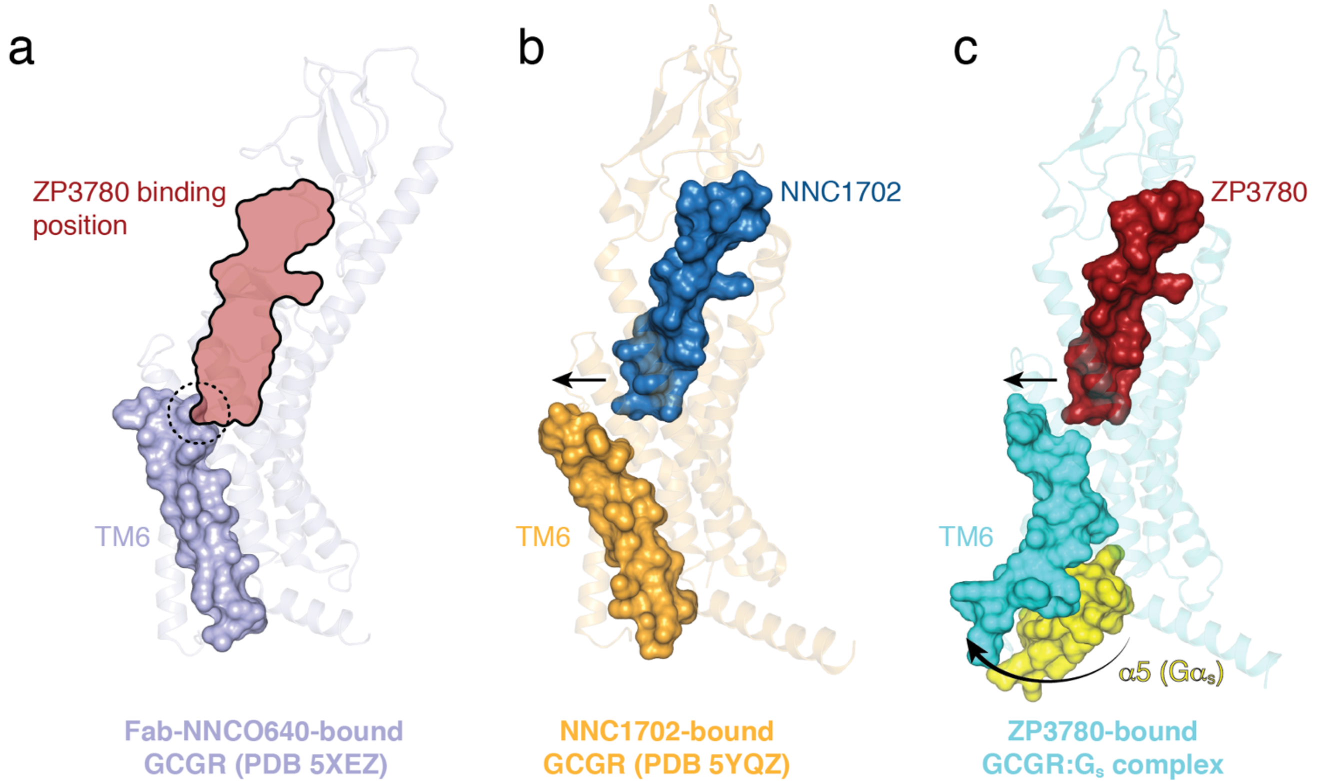


**Supplemental Fig. 6: Peptide and G protein-induced conformational changes in transmembrane domain 6 of GCGR.** a, The binding position of ZP3780 (black outline based on the ZP3780-bound GCGR:$G_s$ complex structure (c)) overlaps with the C-terminal end of TM6 (highlighted with a dashed circle) in the Fab-bound inactive structure of GCGR (PDB 5XEZ, Fab and NNCO640 are not shown for clarity)[3]. b, The extracellular tip of TM6 (orange) moved outward to accommodate NNC1702 (blue) binding in the partial agonist-bound GCGR structure (PDB 5YQZ)[2]. c, In the GCGR:$G_s$ complex structure (cyan), the N- and C- termini of TM6 move away from the receptor core to accommodate ZP3780 (red) and the C-terminal α5 helix of $G\alpha_s$ (yellow), respectively. The necessity for the outward movement of both the extracellular and intracellular sides of TM6 to allow binding of the bulky peptide and G protein might result in the extreme kink formation in class B GPCRs.

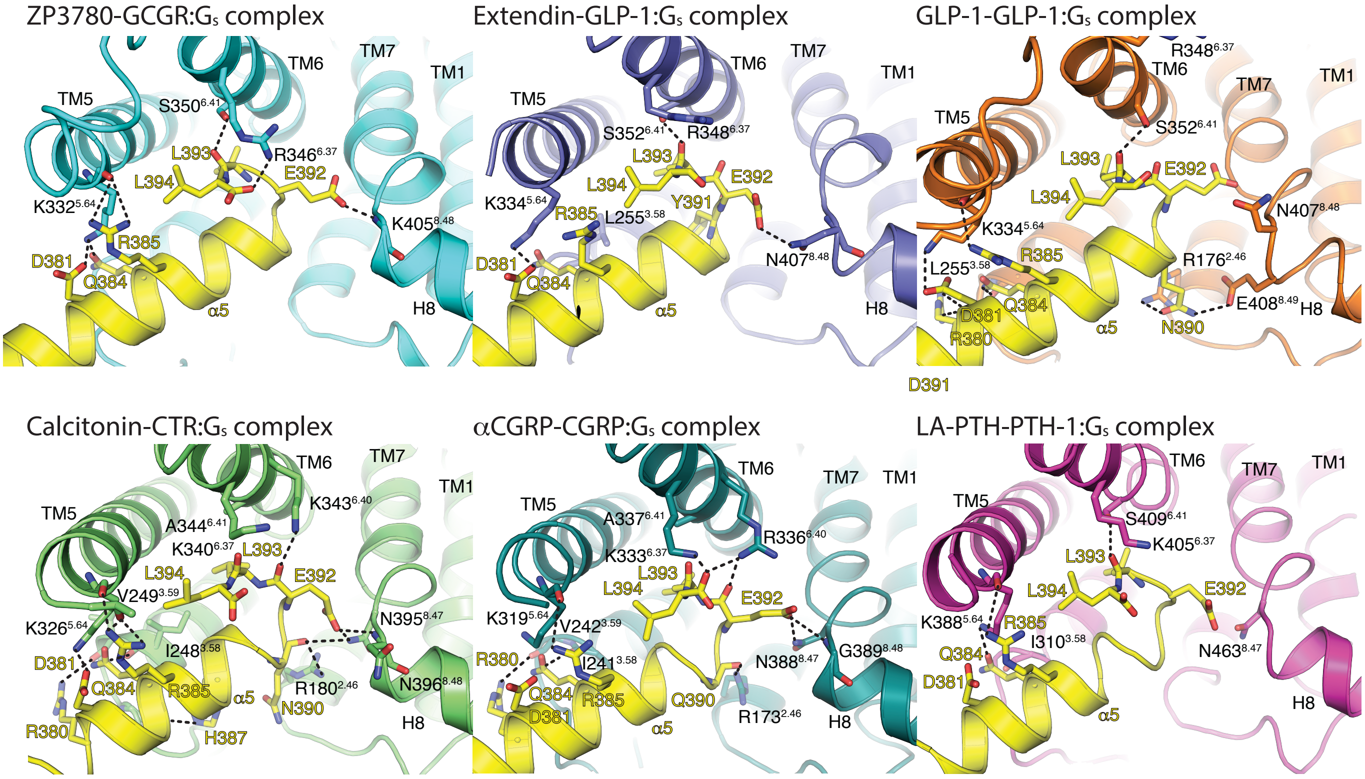


**Supplemental Fig. 7: Comparison of the C-terminal α5 helix binding mode of $G_s$ in Class B GPCRs.** Cytoplasmic views of the GCGR (cyan) with the C-terminal α5 helix of $G\alpha_s$ (yellow), compared to the Extendin-GLP-1:$G_s$ complex (purple, PDB 6B3J)[4], GLP-1-GLP1:$G_s$ complex (orange, PDB 5VAI)[5], Calcitonin-CTR:$G_s$ complex (green, PDB 6NIY)[6], αCGRP-CGRP:$G_s$ complex (deepteal, PDB 6E3Y)[7], and the LA-PTH-PTH1:$G_s$ complex (pink, PDB 6NBF)[8]. H-bonds are shown as black dashed lines.

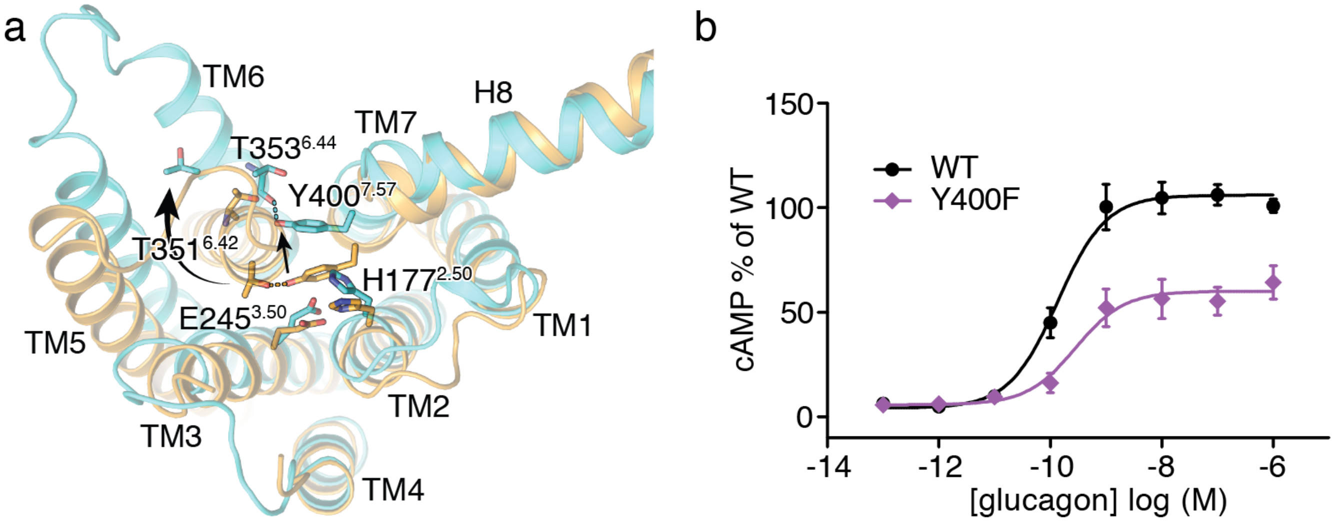


**Supplemental Fig. 8: Rearrangements in the HETx motif of GCGR are important for receptor activation.** a, The TM6 outward movement in the GCGR:$G_s$ complex (cyan) is stabilised by the structural rearrangement in the HETx motif that interacts with the displaced TM6 (viewed from the intracellular side of the receptor). The NNC1702-bound GCGR structure (orange, PDB 5YQZ)[2] is shown for comparison. b, Deletion of the hydroxy group of Y400F$^{7.57}$ in the HETx motif leads to a 2-fold reduction of the GCGR-mediated cAMP signaling. For b, data represent mean ± s.e.m. from at least three independent experiments, performed in triplicates.

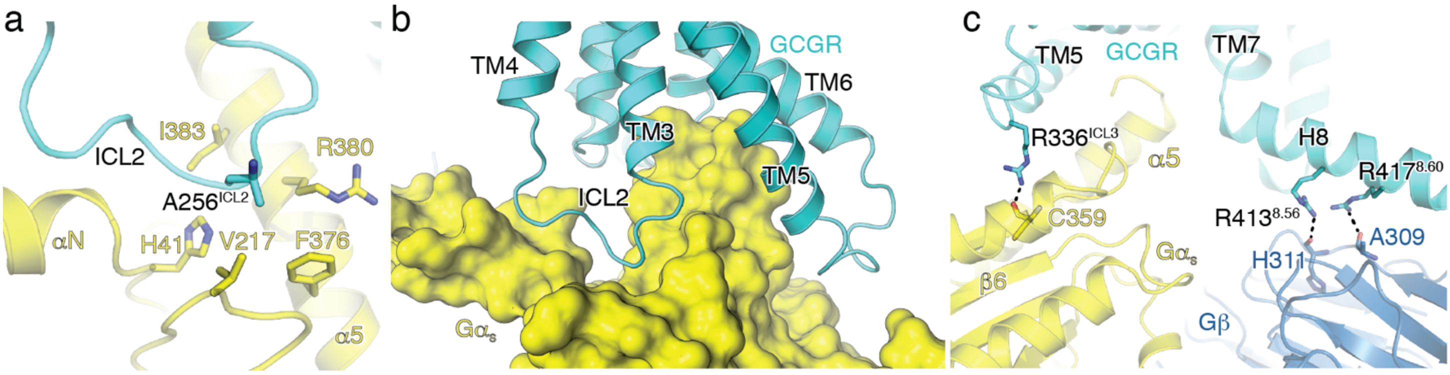


**Supplemental Fig. 9: Interaction of ICL2 and helix 8 (H8) of GCGR with the $G_s$ heterotrimer.** a, ICL2 of GCGR (cyan) forms hydrophobic interactions with residues H41, V217, F376, R380 and I383 of $G\alpha_s$ (yellow). b, ICL2 points into a hydrophobic cavity formed by αN, αN-β1 hinge region and α5 of $G\alpha_s$. c, $R413^{8.56}$ and $R417^{8.60}$ form polar interactions with the backbone atoms of H311 and A309 of Gβ.

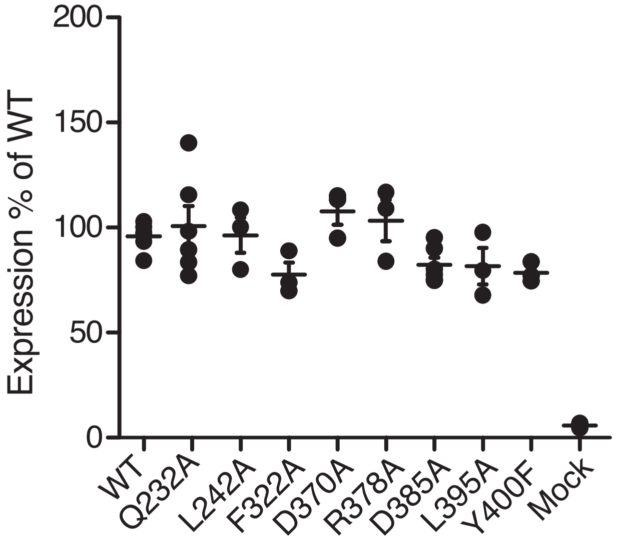


**Supplemental Fig. 10: Surface expression levels of GCGR mutants for functional studies.** For the functional cAMP studies a similar surface expression level of the wild type GCGR and receptor mutants was ensured by titrating the amount of DNA encoding each of the GCGR mutants that was transfected into HEK293 cells. Surface expression levels were determined by cell surface ELISA in 3-6 independent experiments performed in triplicate. The line represent mean ± s.e.m. Oneway ANOVA analysis showed that expression levels were not significantly different (P=0.073, mock not included). Dunnett's Multiple Comparison Test showed that expression levels of each of the mutants were not statistically different from wild type (refer to Suppl. Table 7).

**Supplemental Table 1: Solubility of glucagon and ZP3780 at different pH values.**

| Buffer | Glucagon | ZP3780 |
|---|---|---|
| Acetate pH 4 | >1 mg/ml | >1 mg/ml |
| Acetate pH 5 | >1 mg/ml | >1 mg/ml |
| Phosphate pH 6 | <0.5 mg/ml | >1 mg/ml |
| Phosphate pH 7 | <0.5 mg/ml | >1 mg/ml |
| Phosphate pH 7.5 | <0.5 mg/ml | >5 mg/ml |
| Tris pH 7.5 | <0.5 mg/ml | >1 mg/ml |
| Tris pH 8 | Not tested | >1 mg/ml |
| Glycine pH 9 | Not tested | >1 mg/ml |

**Supplemental Table 2: Comparison of the fibrillation of glucagon and ZP3780**.

| | **Fibrillation** | |
|---|---|---|
| | **Aggregation lag time by ThT** | |
| | **Glucagon** | **ZP3780** |
| | pH 2.5 | pH 7.5 |
| No Agitation | 13.1 ± 1.0 hr | No fibrillation |
| Agitation | 1.8 ± 0.2 hr | 19.6 ± 0.5 hr |

The fibrillation tendency was investigated using the fluorescent probe Thioflavin T (ThT), which detects the presence of amyloid fibrils.
The aggregation propensity was accelerated using high temperature (40°C) combined with or without agitation.

**Supplemental Table 3: Comparison of the binding affinity of glucagon and ZP3780.**

| | **$K_i$ (nM)** | **$pK_i$ ± SEM** | **p-value** | **n** |
|---|---|---|---|---|
| Glucagon | 1.7 | 8.76 ± 0.06 | | 4 |
| ZP3780 | 6.9 | 8.16 ± 0.04 | <0.0001 | 4 |

$pK_i$ values were determined by the Cheng-Prussov equation after non-linear regression of data to single binding site model.
Average Ki-values was calculated from the $pK_i$.
Data are given as mean ± s.e.m. from 4 independent experiments performed in duplicate or triplicate. Statistics was performed as an extra sum-of-squares F test.

**Supplemental Table 4: Comparsion of the EC50 and Emax values of glucagon an ZP3780 in GCGR-mediated cAMP signalling assays.**

| | **EC50 (nM)** | **pEC50 ± SEM** | **p-value** | **Emax (nM cAMP) ± SEM** | **p-value** | **n** |
|---|---|---|---|---|---|---|
| Glucagon | 0.23 | 9.64 ± 0.03 | | 23.4 ± 0.8 | | 4 |
| ZP3780 | 0.33 | 9.49± 0.06 | 0.2237 | 23.0 ± 1.6 | 0.5929 | 4 |

pEC50 and Emax values were determined by fitting data to a four-parameter dose-response curve.
Hill slopes were 1.0 and 1.2 for glucagon and ZP3780, respectively, and not statistically different (extra sum-of-squares F test, p=0.7892).
Average EC50-value was calculated from the pEC50. Emax was measured as nM cAMP pr. well.
Data are given as mean ± s.e.m. from 4 independent experiments performed in duplicate.
Statistics were performed as extra sum-of-squares F tests.

**Supplemental Table 5: Data collection, model refinement and validation.**

| | |
|---|---|
| **Data Collection** | |
| Voltage (kX) | 300 |
| Magnification | 47,169 |
| Total electron dose ($e^-/Å^2$) | 50 |
| Defocus range (µm) | 1.2-2.2 |
| Calibrated pixel size (Å) | 1.06 |
| Micrograph collected (no.) | 3,724 |
| **Data processing** | |
| Extracted particles (no.) | 2,039,910 |
| Particles used for final reconstruction (no.) | 296,516 |
| Final map resolution (Å, 0.143 FSC) | 3.1 |
| Map resolution range (Å) | 2.8-3.6 |
| Map sharpening B factor (Å2) | Pre -90, post -30 |
| **Model content** | |
| Initial models used (PBD code) | 5YQZ (GCGR), 5VAI ($G_s$/Nb35) |
| Total number of atoms | 9,262 |
| No. of protein residues | 1,176 |
| No. of ligands | 0 |
| **Model validation** | |
| CC map vs. model (%) | 80 |
| RMSD | |
| Bond lengths (Å) / Bond angles (°) | 0.007/0.942 |
| Ramachandran plot statistics | |
| Most favored (%) | 94.13 |
| Outliers (%) | 0 |
| Rotamer outliers (%) | 0 |
| C-beta deviations | 0 |
| Clash score | 6.18 |

**Supplemental Table 6**: **cAMP accumulation assay for GCGR mutants.**

| GCGR | EC50 (nM) | pEC50 ± SEM | p-value to WT | Emax (% WT) ± SEM | p-value to WT | n |
|---|---|---|---|---|---|---|
| **WT** | 0.14 | 9.87 ± 0.11 | | 106 ± 3 | | 8 |
| **Q232A** | 51 | 7.30 ± 0.07 | <0.0001 | 92 ± 3 | 0.0407 | 6 |
| **L242A** | 3 | 8.56 ± 0.19 | <0.0001 | 77 ± 5 | 0.0006 | 3 |
| **F322A** | 111 | 6.95 ± 0.17 | <0.0001 | 84 ± 8 | 0.0065 | 3 |
| **D370A** | >100 | n.d | n.d | >74 | n.d | 3 |
| **R378A** | n.d | n.d | n.d | n.d | n.d | 3 |
| **D385A** | >100 | n.d | n.d | >69 | n.d | 6 |
| **L395A** | 0.82 | 9.09 ± 0.19 | 0.0036 | 72 ± 4 | <0.0001 | 3 |
| **Y400F** | 0.28 | 9.55 ± 0.22 | 0.4126 (ns) | 60 ± 3 | <0.0001 | 3 |

pEC50 and Emax values were determined by fitting data to a three-parameter dose-response curve. Average EC50-values were calculated from the pEC50. Emax values were normalized to that of WT in each of the independent experiments.
Data are given as mean ± s.e.m. from the indicated number of independent experiments performed in triplicate n.d., not determined, as parameter could not be reliably fitted.
Statistics were performed by one-way ANOVA, followed by Dunnett's multiple comparisons test to the WT. Each p-value was adjusted to account for multiple comparisons.

**Supplemental Table 7**: **Surface expression of GCGR mutants analysed by Cell surface ELISA.**

| GCGR | Surface expr. (% WT) ± SEM | p-value (to WT) | n |
|---|---|---|---|
| **WT** | 96 ± 3 | | 6 |
| **Q232A** | 101 ± 10 | 0.9932 (ns) | 6 |
| **L242A** | 96 ± 8 | >0.9999 (ns) | 3 |
| **F322A** | 78 ± 6 | 0.3072 (ns) | 3 |
| **D370A** | 108 ± 6 | 0.7609 (ns) | 3 |
| **R378A** | 103 ± 10 | 0.9761 (ns) | 3 |
| **D385A** | 82 ± 3 | 0.4097 (ns) | 6 |
| **L395A** | 82 ± 9 | 0.6015 (ns) | 3 |
| **Y400F** | 78 ± 3 | 0.3638 (ns) | 3 |
| **mock** | 6 ± 0.3 | <0.0001 | 6 |

Data are given as mean ± s.e.m. from the indicated number of independent experiments performed in triplicate.
Statistics were performed by one-way ANOVA, followed by Dunnett's multiple comparisons test to the WT. Each p-value was adjusted to account for multiple comparisons.
